# Temperature regimes impact coral assemblages along environmental gradients on lagoonal reefs in Belize

**DOI:** 10.1101/036400

**Authors:** Justin H. Baumann, Joseph E. Townsend, Travis A. Courtney, Hannah E. Aichelman, Sarah W. Davies, Fernando P. Lima, Karl D. Castillo

## Abstract

Coral reefs are increasingly threatened by global and local anthropogenic stressors such as rising seawater temperature, nutrient enrichment, sedimentation, and overfishing. Although many studies have investigated the impacts of local and global stressors on coral reefs, we still do not fully understand how these stressors influence coral community structure, particularly across environmental gradients on a reef system. Here, we investigate coral community composition across three different temperature and productivity regimes along a nearshore-offshore gradient on lagoonal reefs of the Belize Mesoamerican Barrier Reef System (MBRS). A novel metric was developed using ultra-high-resolution satellite-derived estimates of sea surface temperatures (SST) to classify reefs as exposed to low (low_TP_), moderate (mod_TP_), or high (high_TP_) temperature parameters over 10 years (2003 to 2012). Coral species richness, abundance, diversity, density, and percent cover were lower at high_TP_ sites relative to low_TP_ and mod_TP_ sites, but these coral community traits did not differ significantly between low_TP_ and mod_TP_ sites. Analysis of coral life history strategies revealed that high_TP_ sites were dominated by hardy stress tolerant and fast-growing weedy coral species, while low_TP_ and mod_TP_ sites consisted of competitive, generalist, weedy, and stress-tolerant coral species. Satellite-derived estimates of *Chlorophyll-a (chl-a)* were obtained for 13-years (2003-2015) as a proxy for primary production. *Chl-a* concentrations were highest at high_TP_ sites, medial at mod_TP_ sites, and lowest at low_TP_ sites. Notably, thermal parameters correlated better with coral community traits between site types than productivity, suggesting that temperature (specifically number of days above the thermal bleaching threshold) played a greater role in defining coral community structure than productivity on the MBRS. Dominance of weedy and stress-tolerant genera at high_TP_ sites suggests that corals utilizing these two life history strategies may be better suited to cope with warmer oceans and thus may warrant protective status under climate change.

## Introduction

Coral reefs are threatened locally and globally by anthropogenic stressors such as warming induced by increasing greenhouse gas emissions, excessive nutrients from runoff and sewage effluent, overfishing, and habitat destruction [1,2,3]. Of particular concern are increasing greenhouse gas emissions that continue to cause warming of the global oceans [1,4]. This warming trend is especially troubling in the Caribbean Sea, where rates of warming are higher than in many other tropical basins [5], and where coral cover has declined by up to 80% in recent decades [6]. Elevated sea surface temperature (SST) is the major cause of the breakdown of the essential coral-algal symbiosis, which if widespread results in mass coral bleaching [7,8]. In Belize, the 1998 El Niño bleaching event was the most significant bleaching induced mass coral mortality event on lagoonal reefs over the last 3000 years [9]. These large-scale coral bleaching events are projected to increase in frequency and severity as the climate continues to warm [4,10]. In fact, if ocean warming persists, corals in the Caribbean Sea are predicted to bleach biannually within the next 20-30 years [11], with annual bleaching events occurring as soon as 2040 [12]. Caribbean-wide and global-scale bleaching events are predicted to continue unless corals can increase their thermal tolerance at a rate of 0.2-1.0°C per decade [4].

Annual and daily thermal variability have recently been identified as important factors influencing coral thermal tolerance [13,14,15]. Indeed, previous exposure to thermally variable environments increases a coral’s tolerance to future temperature stress [14,16,17,18], and research suggests that Pacific and Red Sea corals living in areas with high summer maximum SST are less susceptible to bleaching [19,20]. Along the Belize Mesoamerican Barrier Reef System (MBRS) and on Pacific Atolls, corals historically exposed to less thermal variability exhibited slower growth rates and/or greater susceptibility to bleaching in response to SST increases [17,18]. In the Florida Keys, coral growth rates and coral cover were higher in nearshore environments exposed to more variable seawater temperatures than on deeper reefs experiencing more stable temperatures [21]. In contrast, while many studies suggest that high temperature variability leads to higher coral resilience [14,15,16], there is also evidence that corals experiencing moderate long term temperature variability (either annual or daily variation) are better able to cope with stress [13]. Collectively, these studies emphasize the importance of thermal variability on the response of corals to environmental stress, and highlight its capacity to shape coral community composition across a reef system.

Multi-species coral assemblages have recently been proposed to comprise four major life history guilds: competitive (large, fast growing, broadcast spawning, e.g., Caribbean *Acropora spp*.), weedy (small, opportunistic colonizers of recently disturbed habitat, e.g., Caribbean *Porites spp*.), stress-tolerant (massive, slow growing, broadcast spawning, e.g., *Siderastrea siderea*), and generalist (share traits characteristic of all three other groups, e.g., *Orbicella spp*.) [22]. Grouping species by life history strategy allows for prediction of responses to disturbance (e.g., temperature stress) as life history strategies are trait based [23]. Additionally, each guild is expected to be differentially impacted by stressors and life histories predict coral community response to multiple stressors [24]. Therefore, life history strategies offer a more elegant and predictive alternative to traditional genus or species level analysis.

Competitive corals are by definition not very stress tolerant [22]. As such, region-wide decline of these species would be expected as the impact of anthropogenic stressors increase (including coral disease). This decline has already occurred in the Caribbean [6]. Generalist corals became dominant on Caribbean reefs in the late 1970s following mass die off of competitive corals. Generalists are more stress tolerant than competitive species but bleaching and other stressors have led to high mortality of *Orbicella spp*. in the Caribbean [25] and continued decline is expected as temperature stress increases [6,26,27], leading to a decline in reef complexity [28]

Weedy and stress tolerant corals have been shown to be more resilient than competitive and generalist species [22,24], and are hypothesized to dominate warmer and more impacted reefs (e.g., reefs closer to the shore). A shift from dominance of competitive and generalist species to weedy and stress tolerant species occurred on Okinawan reefs following the 1998 El Niño bleaching event [29,30] and an overall decline in coral cover and abundance currently occurring in the Caribbean has been coupled with an increase in abundance of weedy species [27,31]. Interestingly, fossil assemblages from excavated pits on reefs in Panama reveal that mortality and changes in reef communities caused by anthropogenic impact (such as land clearing and overfishing) predate mass bleaching events, indicating that other sub-lethal stressors can impact coral community structure [32,33,34]. Collectively, evidence suggests that differential responses between coral species to increasing anthropogenic stressors may lead to community scale shifts in reef composition from dominance of competitive and generalist species to dominance of stress tolerant and weedy species.

The purpose of the current study was to investigate the impact of thermal regimes on present day coral community composition (coral abundance, species richness, diversity, percent cover, density, and life history strategies) of lagoonal reefs (i.e., region extending from the barrier reef’s crest to the mainland) across the Belize MBRS. A novel GIS-based metric was developed to characterize lagoonal reefs across this reef system into three thermally distinct regimes. Within these three regimes, thirteen reef sites were identified and benthic surveys were conducted to quantify coral community composition. These thermal regimes exist along a nearshore-offshore productivity gradient, which may also influence coral community structure. Quantifying coral community differences among these thermally distinct reefs will help us better predict how coral community structure may be impacted by climate change. Identifying which areas and species are best able to cope with environmental stress (and which are least able) may allow for more targeted management strategies, as it is important to protect both high-risk and low-risk reef sites to improve our chances of conservation success [35].

## Materials and Methods

### Site identification

#### SST Estimate Assembly

Daily 1-km horizontal resolution SST estimates were acquired from the Jet Propulsion Laboratory’s Multi-Scale High Resolution SST (JPL MUR SST) records via the Physical Oceanography Distributed Active Archive Center (PO.DAAC) at the NASA JPL, Pasadena, CA (http://podaac.ipl.nasa.gov). Conventional 1-km resolution satellite-derived SST measurements (infrared, IR) are contaminated by clouds, creating data-void areas. Microwave (MW) data sets can penetrate clouds to gain better temporal coverage, but with a much coarser spatial resolution (25 km) [36]. MUR combines these two datasets to present a more comprehensive and complete SST product. It employs multi-resolution variational analysis (MRBA) as an interpolation method to combine high resolution datasets with more conventional datasets, generating a product that contains no cloud contamination [36]. MUR reports estimates of foundation SST, or SST at the base of the diurnal thermocline (~5-10m depth). Comparison of in-situ temperature (recorded by HOBO^®^ v2 data loggers), MUR, and other SST products revealed that MUR outperforms other products in estimating in-situ temperature, although it also underestimates the temperature corals experience at depth (S1 Fig). However, due to its temporal coverage and temporal resolution, high spatial resolution, lack of cloud contamination, and smaller method error compared to similar products such as Group for High Resolution SST (GHRSST), MUR was determined to be the ideal SST product for use in the current study.

#### Site Classification

Multiple thermal parameters were calculated at different temporal resolutions and examined across thirteen lagoonal reef sites (S1 Table). Lagoonal reefs are located between the barrier reef’s crest and the mainland, and therefore do not include the seaward facing fore-reef. Instead, lagoonal reefs include nearshore reefs, patch reefs, and the back reef. Four thermal parameters produced distinct environments for the reef sites across the Belize MBRS: average annual maximum temperature (S2A Fig), average annual temperature range (S2B Fig), average annual number of days above the regional bleaching threshold of 29.7°C [9] (S2C Fig), and average annual consecutive days above the regional bleaching threshold (i.e., longest potential thermal stress events) (S2D Fig). A metric that combined all four thermal parameters was generated using ArcGIS^©^ in order to assess thermal environments across the Belize MBRS. Data from each of the four parameters in the metric (Table 1) were divided into 8-10 bins (0.5 standard deviations (SD) of the mean) and overlaid on a map of the Belize MBRS. Reefs were not present in areas where the value of any single variable was <1 SD below or >2 SD above the mean (across the entire data set from 2003-2012). For all four parameters, areas that were classified in bins ≥1 SD above the mean were designated high temperature parameter (high_TP_) sites (Fig 1). Moderate temperature parameter (mod_TP_) sites were classified as areas where all values were 0.5 to 1 SD above the average annual temperature range and the average annual maximum temperature, and within 1 SD of the average annual consecutive days and the average annual number of days above the regional bleaching threshold (Fig 1). Low temperature parameter (lOW_TP_) sites were classified as bins that were 0.5 SD above the average to 2 SD below the average for annual temperature range and annual maximum temperature, and below the average for consecutive and annual days above the regional bleaching threshold (Fig 1). Using the metric presented in Fig 1, fifteen sites were identified, thirteen of which were visited and surveyed in November 2014 (the two northernmost high_TP_ sites were not surveyed as corals were not located within the marked geographic area) (Table 1, Fig 1).

**Fig. 1.**
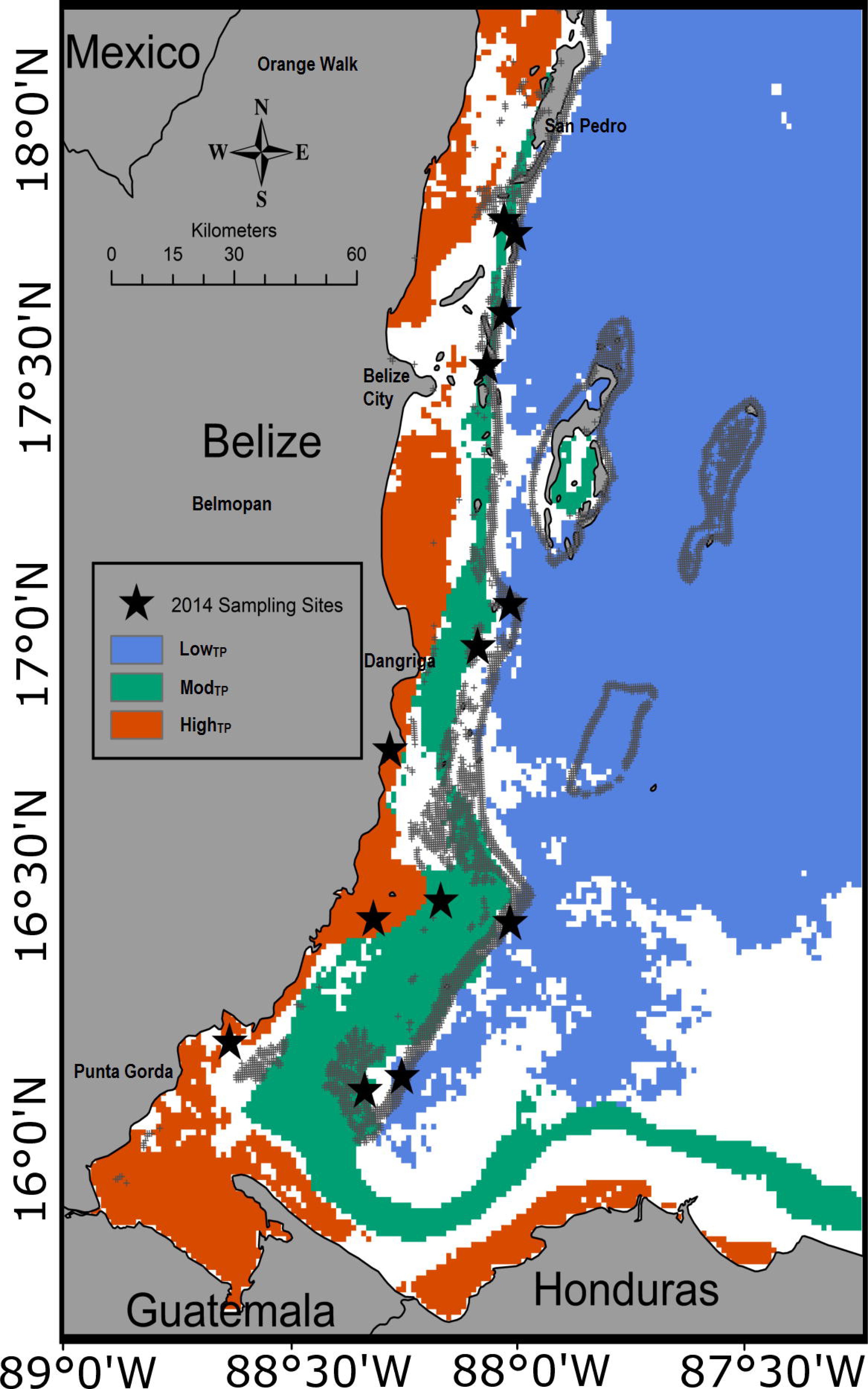
Thermal Regimes and Site Locations.

The Belize Mesoamerican Barrier Reef System (MBRS) classified by site type based on four thermal parameters. Blue, green and red regions represent low_TP_, mod_TP_, and high_TP_ areas across the reef system. Stars indicate surveyed sampling sites.

**Table. 1.** Thermal Parameters Used For Site Classification.

| Factor | Min | Mean | Max | SD | low<sub>TP</sub> Sites | mod<sub>TP</sub> Sites | high<sub>TP</sub> Sites |
| --- | --- | --- | --- | --- | --- | --- | --- |
| Mean Annual Max Temp | 30.2°C | 30.6°C | 31.3°C | 0.27°C | 30.2-30.8°C | 30.8-30.9°C | 30.9-31.3°C |
| Mean Annual Temp Range | 4.4°C | 5.2°C | 7.1°C | 0.69°C | 4.4-5.5°C | 5.5-5.9°C | 5.9-7.1°C |
| Mean Annual Days Above Bleaching Threshold | 20.0 days | 40.1 days | 78.4 days | 14.3 days | 20.0-40.1 days | 40.1-54.4 days | 54.4-78.4 days |
| Mean Consecutive Days Above Bleaching Threshold | 3.0 days | 4.8 days | 7.5 days | 0.92 days | 3.0-4.8 days | 4.8-5.7 days | 5.7-7.5 days |

Values for the four thermal parameters included in site selection metrics. Values are all averages from 2003-2012 and include measurements for minimum, mean, maximum, and standard deviation (SD) for each thermal parameter. The range at which each factor was classified as low_TP_, mod_TP_, or high_TP_ site is also shown.

### Benthic surveys

In November 2014, benthic surveys were performed at the thirteen reef sites. Depth of each reef site was standardized to 3-5m. Reef types surveyed included back reefs, patch reefs, and nearshore reefs. A team of three divers surveyed six belt transects (dimension 6 × 10 m) at each site following Atlantic and Gulf Rapid Reef Assessment (AGRRA) methodology [37]. Briefly, a diver classified the genus and species of every coral >6cm^2^ within 1m of the transect line along a 10m transect. The number and size (length, width, and height) of individual colonies of each coral species were recorded on underwater data sheets. The collected data were used to calculate coral species diversity, abundance, richness, and coral life history (following Darling *et al*. [22]) for each site.

Additionally, six video belt transects (1 × 20m) were also performed at each site using GoPro^®^ cameras attached to PVC stabilizing apparatuses allowing each diver to stabilize the camera while surveying transects. Video transects were analyzed at the University of North Carolina at Chapel Hill (UNC-Chapel Hill) in a manner similar to the AGRRA method used in the field, except two additional parameters (percent coral cover and coral density) were calculated. Results of the diver and video transect surveys were not significantly different (*p*=0.300). As a result diver and video survey data were pooled at each site when possible. Full details and a comparison of the methods employed are available in S1 Appendix.

### Coral life history

Coral species were grouped into four life history strategies as previously described by Darling *et al*. 2012 [22]. In their study, Darling *et al*. 2012 identified four life history guilds for corals based on multivariate trait analysis: competitive, weedy, stress-tolerant, and generalist [22]. The four guilds are primarily separated by colony morphology, growth rate, and reproductive rate. The classification was based on a thorough sampling of global Scleractinian coral diversity. Each coral that is included in a guild in Darling *et al*. 2012 [22] was classified into the appropriate guild for this study and comparisons of life history strategies between sites and site types were made.

### Chlorophyll-a

Eight-day composite 4-km horizontal resolution *chlorophyll-a* (*chl-a*) estimates over the interval 2003-2015 were obtained from NASA’s Moderate Resolution Imaging Spectroradiometer (AQUA MODIS) via NOAA’s Environmental Research Division’s Data Access Program (ERDDAP) [38]. Eight-day composite data were selected in order to minimize gaps in data from cloud cover. Unlike the MUR SST data used for temperature calculations, there is no integrated, high-resolution product for *chl-a*. Similar to temperature calculations, monthly and yearly average *chl-a* values were calculated for each survey site (S2E Fig). *Chl-a* is a widely used proxy for primary productivity and nutrient delivery in seawater [39,40], as it is the main photosynthetic pigment present in phytoplankton which can often quickly deplete nutrient concentrations below detectable limits. It has been shown that remotely sensed data, such as *chl-a* concentration, yields better metrics for water quality than traditional measures such as distance from shore and distance from the nearest river [41]. Here, *chl-a* data are used as a proxy for primary production across the Belize MBRS.

### Statistical analysis

Standard deviations used for temperature bins and site classification were calculated in ArcGIS^©^. All other statistical analysis were carried out in R 3.2.2 [42]. Transect averaged survey data for species richness, abundance, Shannon diversity, coral cover, coral density, and log-transformed *chl-a* data were analyzed using analyses of variance (ANOVA). Three fixed factors were included in the ANOVA (survey method, site, and site type) for species richness, abundance, and Shannon diversity. Only two fixed factors (site and site type) were included in the ANOVA for coral cover and coral density, since only data from video surveys were used to calculate these averages. Two fixed factors (site and site type) were included in the ANOVA for *chl-a* concentrations since they were calculated using satellite estimates and survey type was not a factor.

If factors were significant (*p*<0.050), a post-hoc Tukey’s HSD test was used to evaluate the significance of each pair-wise comparison. Spatial autocorrelation was evaluated using Moran’s I [43]. Significant *p-values* for Moran’s I (*p*<0.050) indicate an effect of spatial autocorrelation. Spatial autocorrelation was only a factor for coral cover (*p*=0.040). To correct for the effect of spatial autocorrelation, the cut-off value for significance within the ANOVA for coral cover was decreased to *p*<0.010, following Dale and Fortin [44].

To visualize coral community differences between site types, non-metric multidimensional scaling (NMDS) ordination was implemented using Bray-Curtis similarity coefficients in the vegan package in R [45]. An optimal stress test was performed to determine the optimal k value (k=20). Resulting NMDS scores were visualized in two-dimensional ordination space. A PERMANOVA test was performed to analyze the site type differences using the *adonis* function in the vegan package in R [45].

Linear models tested for the influence of temperature parameters and *chl-a* on the variation observed along NMDS1 and NMDS2 (within and between site type community variations). Linear models were run using the *lm* function in R (R Core Team, 2014). R^2^ and *p*-values were calculated for each parameter based on each linear model (S2 Table). For NMDS1, data were also divided by site type in order to assess within site type variation (S3 Table).

### Ethics statement

All research related to this projected was completed under official permit from the Belize Fisheries Department (#000045-14).

## Results

### Coral community composition

Combined results of AGRRA diver surveys and GoPro^®^ video surveys for all thirteen sites revealed that coral species richness varied as a function of site location (*p*<0.001) as well as site type (*p*=0.002). Coral abundance was significantly lower at highTP sites compared to IOWTP (*p*=0.005) and modTP (*p*=0.020) sites, but was not significantly different between IOWTP and modTP sites (Fig 2A). Coral cover, Shannon diversity, coral density, and species richness also followed these same patterns (*p*≤0.020; Fig 2B-E). NMDS analysis of the ecological parameters showed that community structure was significantly different (stress=0.018, adonis test *p*=0.006) between high_TP_ sites and low_Tp_/mod_TP_ sites along the NMDS2 axis, but was not different between low_TP_ and mod_TP_ sites (*p*>0.050) (Fig 3). The most dominant taxa at IOWTP and modTP sites were *Orbicella spp, Porites spp, Undaria spp, S. siderea*, and *Pseudodiploria spp*, while at high_TP_ sites they were *Siderastrea spp., P. astreiodes*, and *Pseudodiploria spp*. Variation along the NMDS1 axis represents within site type differences while variation along the NMDS2 axis represent between site type differences (Fig 3).

**Fig. 2.**
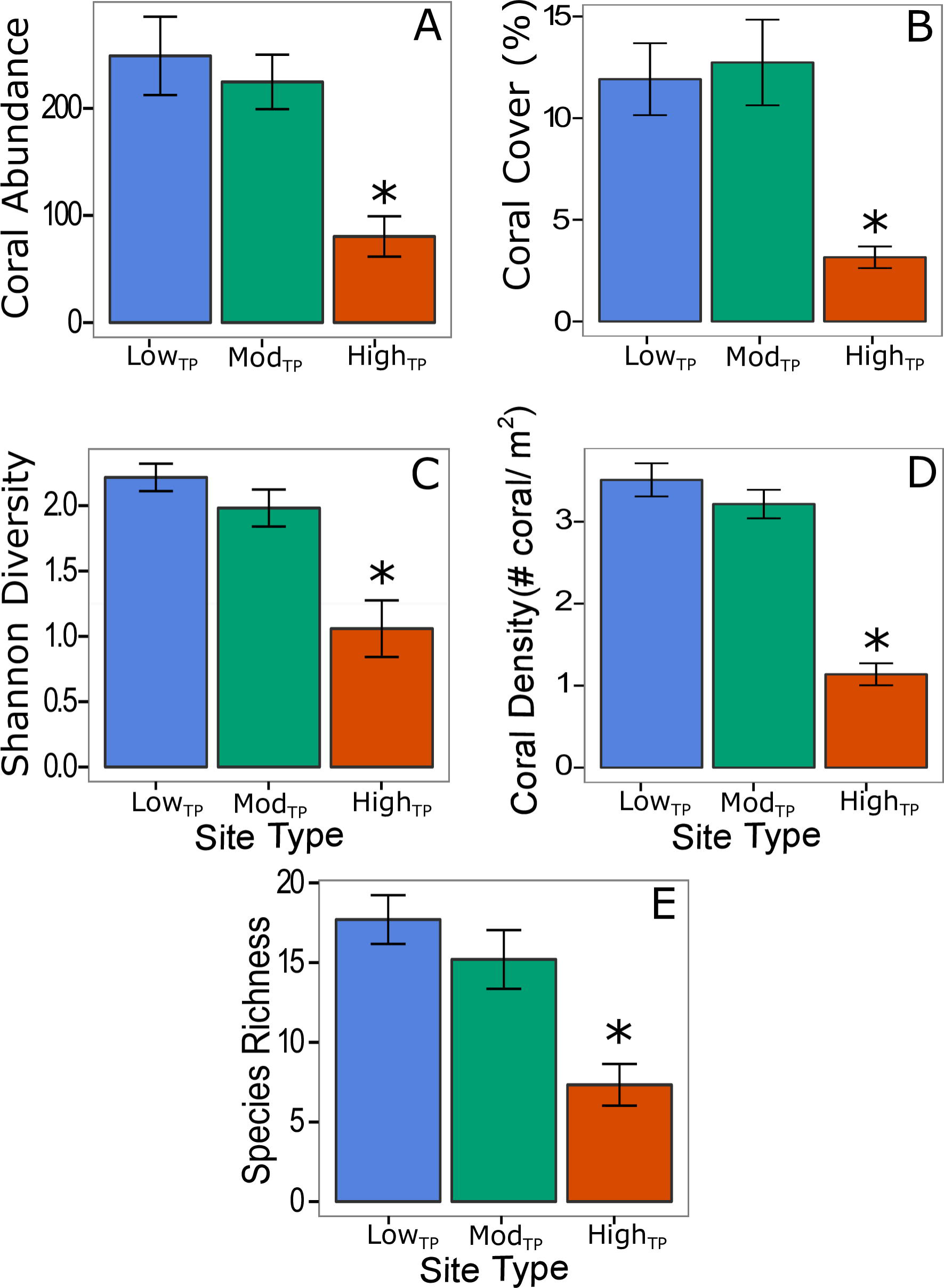
Differences in coral community structure across site type.

Average coral abundance (A), percent coral cover (B), coral species diversity (C), coral density (D), and coral species richness (E) at each site type. Statistically significant differences (*p*<0.05) are marked with an *. Blue, green, and red bars (± 1 SE) represent IOWTP, modTP, and highTP, respectively.

**Fig. 3.**
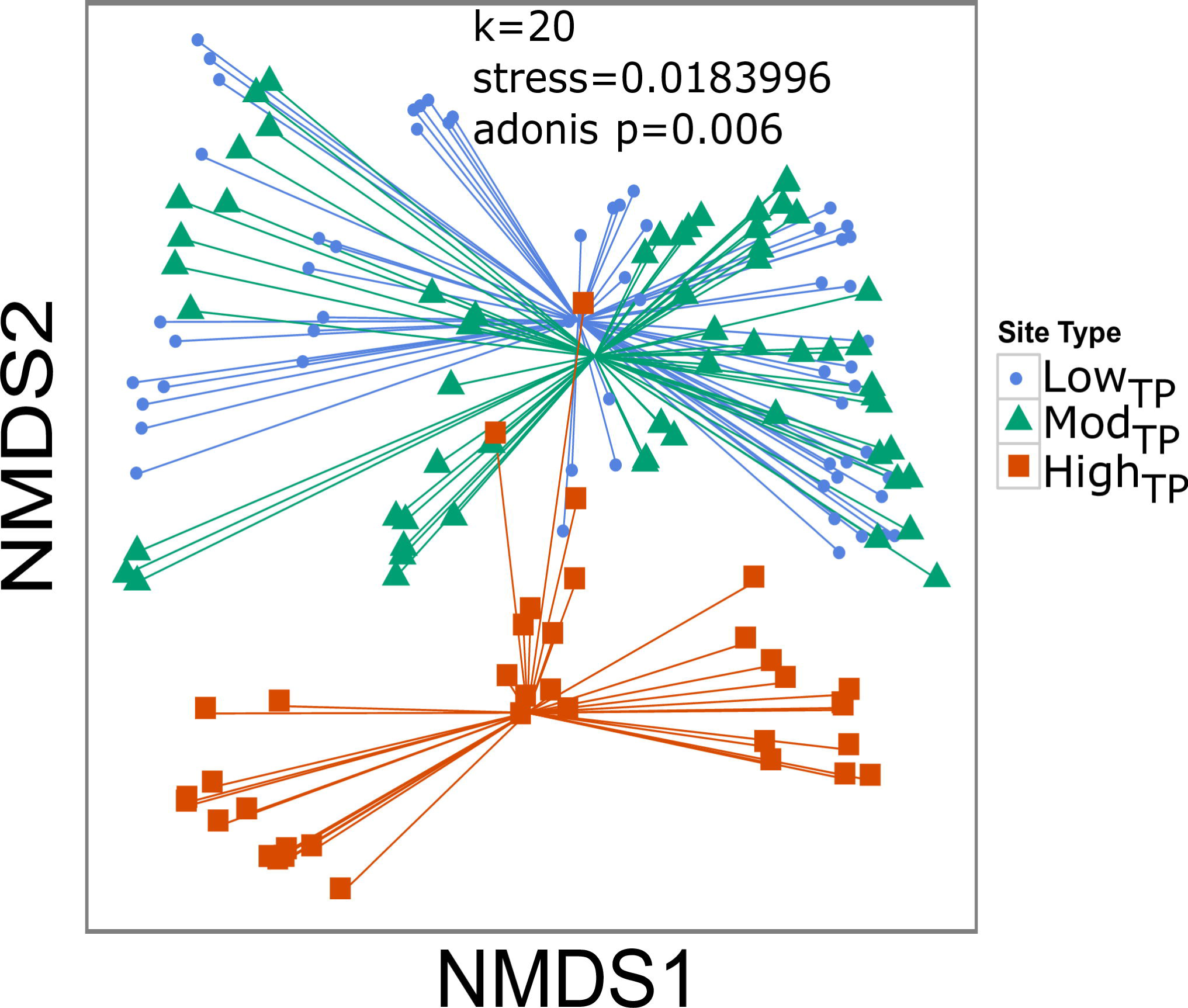
NMDS of coral community variables by site type.

Nonmetric multidimensional scaling (NMDS) plot of coral community differences clustered by site type. Blue circles, green triangles, and red squares represent IOWTP, modTP, and highTP site types, respectively.

Linear modeling of temperature and productivity parameters against NMDS1 and NMDS2 revealed that average annual maximum temperature, average annual temperature range, average annual days above the bleaching threshold, and average annual consecutive days above the bleaching threshold all had significant effects on the NMDS1 variation. All four temperature parameters, as well as *chl-a*, also had significant effects on NMDS2 variation (S2 Table; S3 Fig). Average annual consecutive days above the bleaching threshold explained the most variation for NMDS1 and NMDS2 (R^2^=0.1026, 0.604 respectively; *p* <0.001 for both; S2 Table; S3 Fig).

Linear regressions of temperature parameters and *chl-a* within site types along NMDS1 revealed significant effects (*p*<0.050) of average annual maximum temperature, average annual days above the bleaching threshold, and average annual consecutive days above the bleaching threshold for all site types, average annual temperature range for modTP and high_TP_ sites, and *chl-a* for high_TP_ sites only (S3 Table; S3 Fig). Average annual days above the bleaching threshold yielded the highest R^2^ for lOW_TP_ and mod_TP_ sites, while average annual temperature range yielded the highest R^2^ for high_TP_ sites (S3 Table; S3 Fig).

### Coral life history

Site exhibited a significant effect on the number of corals in each of the four coral life history guilds [22] (*p*<0.001). The distribution of coral life history strategies differed significantly between low_TP_ and high_TP_ site types (*p*=0.049; Fig 4), while mod_TP_ sites did not differ from low_TP_ or high_TP_ sites (Fig 4). Overall, there appears to be a pattern of lower abundances of all life history guilds at highTp sites compared to low_TP_ sites. Competitive species were not present and generalist species were only present in very small number at high_TP_ sites.

**Fig. 4.**
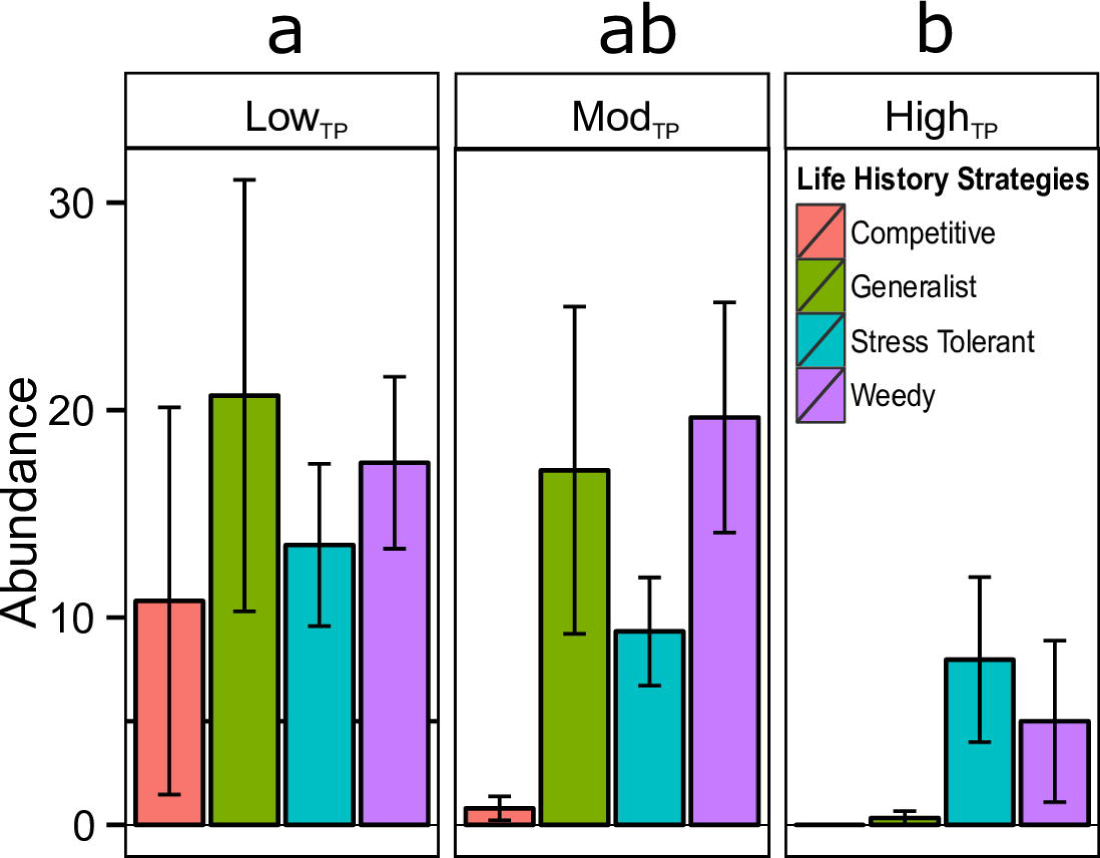
Coral life history strategy by site type.

Abundance (count) of corals (±1 SE) grouped by life history (from Darling *et al*. 2012). Letters ‘a’ and ‘b’ show significant differences between site types (*p*<0.050) acquired from post hoc Tukey tests.

### Chlorophyll-a

Annual average *chl-a* concentrations varied over time and differed by site type (*p*<0.001), but were consistently lowest at low_TP_ sites and highest at high_TP_ sites regardless of year (Fig 5A). *Chl-a* concentrations averaged over 2003-2015 were significantly different across all three site types (*p*<0.001 in all cases). Low_TP_ sites exhibited the lowest average 13-year *chl-a* concentrations. Mod_TP_ sites exhibited average 13-year *chl-a* concentrations that were significantly higher than low_TP_ sites, but significantly lower than high_TP_ sites. HighTP sites exhibited significantly higher average 13-year *chl-a* values than both low_TP_ and mod_TP_ sites (*p*<0.001 in all cases, Fig 5B). The pattern seen in *chl-a* concentrations is positively correlated with the patterns seen in all temperature parameters (*chl-a* and temperature parameters are lowest at low_TP_ sites and highest at high_TP_ sites) (Fig 1, S2 Fig).

**Fig. 5.**
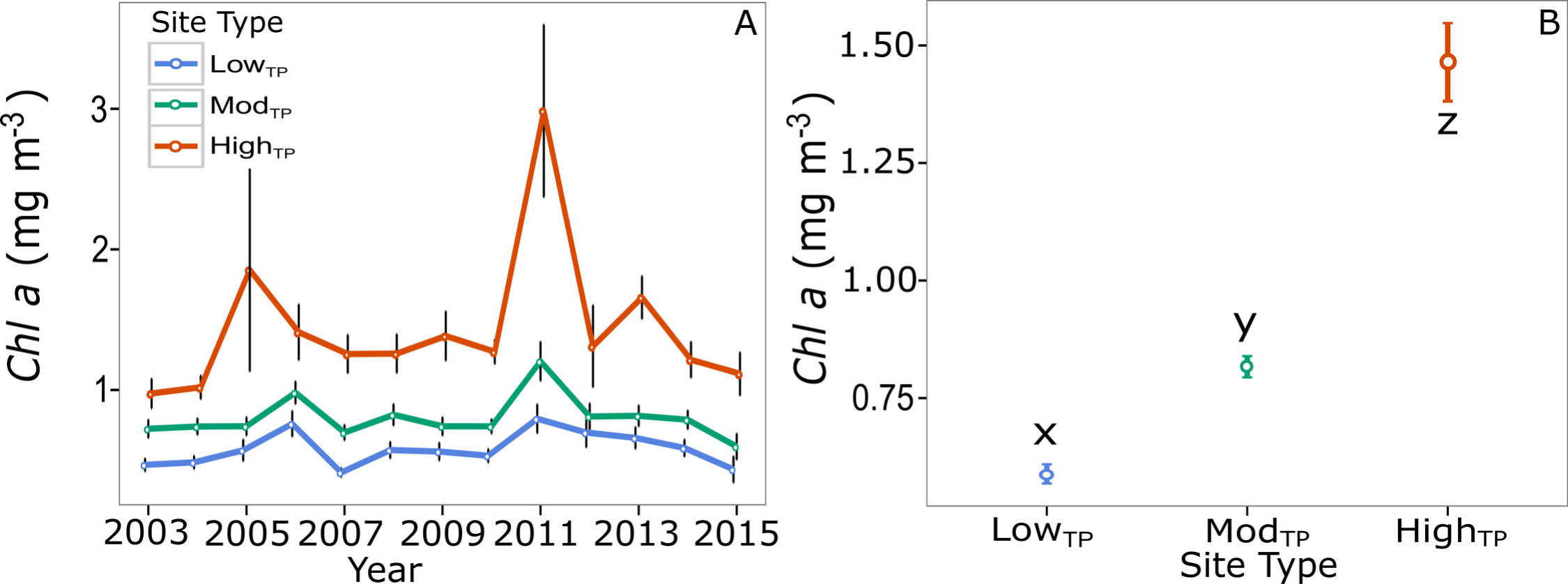
Average *chl-a* by site type.

*Chl-a* concentration by site type (±SE) Annual average *chl-a* for low_TP_ (blue), mod_TP_ (green), high_TP_ (red) site types over the interval 2003-2013 (A). *Chl-a* concentrations averaged over the 13-year interval (B). Letters x, y, and z indicate results of post hoc Tukey tests showing significant differences in 13-year *chl-a* concentrations across site types (*p*<0.050).

## Discussion

### Coral community composition

Coral species richness, abundance, diversity, density, and percent cover were all lower at high_TP_ sites compared to low_TP_ and modip sites (Fig 2). Differences in coral community composition between high_TP_ sites and low_TP_/mod_TP_ sites are historically explained by more stressful conditions nearshore and less stressful conditions offshore [46,47]. These nearshore stressors include, but are not limited to temperature, eutrophication, sedimentation, and wave energy [46,47]. Our findings suggest that lower coral species richness, diversity, abundance, percent cover, and density at high_TP_ sites may be driven by high thermal variability, elevated maximum temperatures, and prolonged duration of exposure to temperatures above the bleaching threshold; three variables that have been shown to cause coral community decline [13,29,48,49,50]. These temperature parameters were more strongly correlated with changes in coral community composition between site types than with *chl-a* (S3 Fig), indicating that they likely play a greater role in determining coral community composition than productivity. High weekly thermal variability has also been shown to correlate with low coral cover on nearshore reefs in the Florida Keys [13]. Therefore, differences in thermal variability observed across site types may have influenced coral community composition in Belize.

Our findings are contrary to the results of Soto *et al*. (2011) [13], which showed that reef sites with moderate temperature variability (equivalent to mod_TP_ sites in the current study) in Florida had higher coral cover than sites exposed to low (offshore deep reefs) or high temperature variability. Soto *et al*. (2011) [13] suggests that corals exposed to moderate weekly thermal variation are able acclimatize to a wide range of environmental conditions, making them more resilient than corals that experience less variation. At the same time, corals exposed to extremely high thermal variation generally do not survive[13]. Our results may contrast with that of Soto et al. (2011) because fore reef locations were not included in the present study (i.e., low_TP_ sites are located in the back reef). Our high_TP_ sites follow the same pattern seen in Soto *et al*. (2011) [13] as they have lower coral cover than mod_TP_ sites (Fig 2).

Our results also contrast those of Lirman and Fong (2007) [21], which showed that nearshore reefs (equivalent to our high_TP_ sites) exhibited higher coral cover and growth rates than offshore reefs (equivalent to our low_TP_ sites) in the Florida Keys. Interestingly, these nearshore Florida reefs also experienced lower water quality than the offshore reefs [21]. The authors hypothesized that higher coral cover and growth rates on nearshore reefs were due to the ability of some corals to switch trophic mode under adverse conditions [21], a pattern that has been observed in previous studies, but was not quantified in the current study [51,52]. Differences in coral community composition between the Florida Reef tract and the Belize MBRS may explain our contrasting results in coral cover as nearshore patch reefs in Florida appear to have relatively high numbers of *Orbicella spp*. [21], whereas high_TP_ sites in Belize were almost void of this species.

### Life history strategies

In the current study, high_TP_ sites contained no competitive species, few generalists, and were dominated by stress-tolerant and weedy genera, while both low_TP_ sites and mod_TP_ sites contained all 4 life history types (Fig 4). Low_TP_ sites contained all four life history strategies in roughly equal proportions. mod_TP_ sites were similar but with fewer competitive species than low_TP_ sites, and high_TP_ sites had comparatively fewer of all four life histories, but were dominated by weedy and stress tolerant genera. Shifts toward weedy and stress tolerant genera under climate change conditions were predicted by Darling *et al*. (2012) [22], and have been recorded in many areas of the world [29,53], including the Caribbean [25,31,54]. Even in the face of region-wide decline in coral cover and decrease in abundance of competitively dominant species [6], some weedy species, such as *Porites astreoides*, are actually increasing in prevalence within the Caribbean [31]. This weedy coral species is likely able to succeed in high stress environments due to its ability to brood and mature quickly, which allows it to rapidly colonize a recently disturbed area [22,31].

In contrast, a stress-tolerant species such as *S. siderea* is likely able to survive in high_TP_ environments due to its massive size and long life span, which allows it to sustain a population in the absence of successful recruitment. This can increase the long-term survival potential of this species in harsh conditions [55]. These two contrasting strategies seem most effective in high_TP_ environments (Fig 4), and are likely to be most effective in future conditions as the oceans continue to warm. This prediction is corroborated by Loya *et al*. (2001) [29], who showed that mounding (e.g., *S. siderea*) and encrusting (e.g., *P. astreoides*) species survived a mass bleaching event in 1997-1998 better than corals of other morphologies (e.g., branching). Ten years after the bleaching event these same types of coral continued to dominate. However, some branching species recovered and increased in abundance [56]. In the current study, branching species were almost non-existent in high_TP_ sites, which indicates that these sites have experienced a recent thermal stress event or are exposed to chronic stress (e.g., temperature, eutrophication) that prevents such species from succeeding in these environments. It is also possible that high_TP_ sites are more frequently disturbed than both low_TP_ and mod_TP_ sites. Disturbances such as bleaching events, eutrophication, sedimentation, and overfishing are known to cause declines in coral cover, species richness, and diversity [29,30]. These more disturbed or impacted reefs can then become dominated by stress-tolerant corals and corals that quickly colonize areas after a perturbation (i.e., weedy corals) [13,29,30,57], as observed in the current study (Fig 4). Historical and/or geological investigation of reef assemblages (i.e., through pit excavating or coring of reef framework [9,32,34]) would be a useful next step, as it would allow insight into how reef communities within the three thermal regimes have changed after disturbances and over long periods of time.

### Influence of primary productivity on coral community composition

Cross-reef *chl-a* concentrations follow the same patterns as temperature (elevated nearshore and decreasing with increasing distance from the Belize coast) (Fig 1, S2 Fig). This means that reefs with higher *chl-a* concentrations have lower coral species richness, abundance, diversity, density, and percent cover. This supports a previous finding that shows a strong negative relationship between *chl-a* and coral cover, species richness, and abundance at nearshore reefs on the Great Barrier Reef (GBR) [58]. However, our results reveal that *chl-a* concentrations are not strongly correlated (R^2^=0.040) with changes in coral community structure (e.g., percent cover, abundance, diversity, species richness, and density) across site types (S5H Fig), suggesting that *chl-a* concentrations may not best explain differences in community composition between site types in Belize. This may be due to spatial scale (e.g., we focused on nearshore, patch reef, and back reef sites as opposed to exclusively nearshore sites) [58], or the coarse scale of the *chl-a* dataset (4 km × 4 km grid; each survey site is <1 km). Focusing on variation within nearshore (high_TP_) sites, we do see a correlation between *chl-a* and changes in coral community structure (S4H Fig), which supports results from previous work [58,59].

### Other potential factors influencing coral community structure across reef types

#### Eutrophication

Eutrophication has led to local degradation of reefs [60,61,62]. However, larger scale (regional) reef degradation due to nutrients alone has not been quantitatively shown [63]. Wooldridge (2009) [64] demonstrates that lower water quality (e.g., higher nutrient concentrations) are linked to lower bleaching thresholds on nearshore reefs in Australia. If bleaching thresholds are depressed at high_TP_ sites for some species, it may help explain lower diversity measured at these sites, as they experience warmer temperatures and spend more time above the regional bleaching threshold than do mod_TP_ and low_TP_ sites (S2 Fig). While *chl-a* does not correlate well with changes in coral community structure in this study (S3 Fig), it should be noted that *chl-a* is an estimate of nutrient delivery and primary productivity, not a measurement of the concentration of any one nutrient pool. Due to this limitation, manipulative field experiments such as Vega-Thurber et al. (2014)[65] and Zaneveld et al. (2016)[66] are needed to understand the influence of nutrients on coral community structure and bleaching thresholds at local scales.

#### Sedimentation

Coastal (nearshore) reefs throughout Belize are influenced by runoff from smaller local rivers, and reefs in southern Belize experience additional runoff and river plumes originating from larger watersheds in Honduras and Guatemala [67,68]. It has been previously shown that *Orbicella faveolata* corals on reefs with higher sedimentation rates exhibited suppressed skeletal extension rates for a longer duration than corals on reefs with lower sedimentation rates following the 1998 bleaching event in Belize [69]. In contrast, increased sedimentation did not affect skeletal extension of *S. siderea* or *P. astreoides* corals in Puerto Rico [70]. The results of these two studies suggest that there may be species-specific responses to increased sedimentation rates. In Barbados, reefs with high sedimentation rates were dominated by coral species with high recruitment and high natural mortality (e.g., *P. astreoides*) and reefs with lower sedimentation rates were dominated by coral species with lower recruitment and low natural mortality (e.g., boulder corals) [71]. As sedimentation rate was not quantified in this study, the impacts of sedimentation on coral community structure are not clear.

#### Circulation and wave energy

The Belize MBRS lies west of the Honduras Gyre, a hydraulic feature that recirculates water inside the Cayman basin [72]. The coastal waters of northern Belize are influenced by the Cayman and Yucatan currents, which move water northwest up the coastline toward Mexico [72,73,74,75]. In central and southern Belize, current velocities are lower and dominant circulation patterns are less consistent throughout the year [74]. However, currents appear to bring water and potentially pollution, nutrients, or sediment plumes from coastal Honduras and Guatemala west to southern Belize where they recirculate before slowly moving northward [67,68,74,75,76,77,78]. These circulation patterns have the potential to influence the stress tolerance of corals across sites and latitude in the current study. Our results reveal no spatial autocorrelation between sites for any of our measured variables with the exception of *chl-a* suggesting that the influence of these currents may be minimal. Additionally, wave energy may play a role in shaping coral communities. Wave energy may be elevated at low_TP_ sites as they are located near channels in the fore reef and may not be as sheltered by the reef crest as other mod_TP_. Similarly, wave energy may be elevated at high_TP_ sites due to the large fetch between the reef crest and nearshore reefs and the prevailing wind direction from offshore to inshore.

#### Light

Irradiance (light intensity) has been shown to decrease along an offshore-nearshore gradient on the GBR as *chl a* concentrations increase [79]. *Chlorophyll-a* concentrations increase with proximity to shore in Belize (Fig 1), so this pattern of decreasing light intensity towards the nearshore likely holds for Belize as well. However, in southern Belize offshore reefs (and nearshore reefs) are subject to seasonal sedimentation and runoff from larger rivers in Honduras and Guatemala [77,78]. Irradiance is a known stressor, proven to cause coral bleaching alone or in consort with elevated temperatures [80]. Although depth was held constant in the present study, it is possible that differing light levels both between site types and between individual sites may influence coral community composition across the site types investigated in the current study.

#### Proximity to human populations

Declining health of coral reefs worldwide has been linked to land-based stressors including nutrients and human use and exploitation (e.g., overfishing) [60,80,81] as well as proximity to sources of these stressors (e.g., major human population centers) [82]. However, not all reefs that are near to or influenced by land-based stressors are unhealthy [21,83]. Some of the study sites were within close proximity to a major human population center, particularly the high_TP_ sites (populations of major towns and cities in Belize can be seen in S4 Table). Analysis of spatial autocorrelation revealed no significant differences between high_TP_ sites or between high_TP_ sites and sites that were further offshore, suggesting that proximity to human population centers did not have a major impact on coral community composition.

### Conclusions

High_TP_ reefs exhibit lower coral diversity, abundance, species richness, and cover than do low_TP_ and mod_TP_ reefs in Belize. These high_TP_ reefs are exposed to higher annual temperatures, greater temperature variability, more time above the regional bleaching threshold, elevated *chl-a* concentrations, and likely increased sedimentation rates and lower flow than low_TP_ and mod_TP_ reefs. Temperature parameters, most notably time spent above the bleaching threshold, correlate best with differences in coral community structure. In addition, stress-tolerant and weedy coral life history strategies dominate at high_TP_ reefs. Due to exposure to generally more stressful environmental conditions, high_TP_ reefs may offer a snapshot into the projected future of coral reefs as they become increasingly exposed to local (pollution, runoff, land-use change, and overpopulation) and global (warming and acidification) stressors. Previously, such reefs have been suggested as possible refugia against climate change [84]. Globally, this would mean a shift towards dominance of stress-tolerant and weedy corals [53]. Such a shift would dramatically impact the structure and function of reefs, essentially creating novel ecosystems [85]. HighTP reefs should be protected in addition to more pristine reefs in order to improve conservation success [35]. More pristine reefs should be protected as they contain more diversity and provide more ecosystem services than do high_TP_ reefs [86]. However, high_TP_ reefs host coral holobionts that may be best suited to survive in future ocean conditions. To ensure survival and future success of reefs while maintain current diversity both heavily impacted and pristine ecosystems must be protected. The results of the current study highlight the need to better protect and understand impacted nearshore reef systems, including investigations into what conditions allow more sensitive species (e.g., competitive and generalist) to survive and persist on nearshore reefs.

## Data Accessibility

All data are archived on PANGAEA at the following DOI: doi.pangaea.de/10.1594/PANGAEA.859972

## Supporting Information

**S1 Appendix. Additional detail of AGGRA and video survey methods**

**S1 Fig. *In situ* temperature versus satellite SST products**

A comparison of *in situ* temperature and MUR SST. *In situ* loggers were collected from 6 sites along the BBRS (site numbers are listed in the gray headers above each panel). Each panel shows a month by month comparison of *in situ* logger measurements and SST products. Zero on the y-axis represents the average value for the Hobo Pro V2 loggers at each site. Red errors bars the standard deviation over a month for each logger. Gray squares show average values for an additional *in situ* logger that was placed at the site (± 1 standard deviation). Blue, green, and black symbols show monthly average values for various SST products (± 1 standard deviation).

**S2 Fig. Temperature parameter and *chl-a* maps**

Maps showing the 4 parameters used to calculate site type: yearly maximum temperature (A), Mean annual temperature range (B), Annual mean number of days above the bleaching threshold (C),Annual mean consecutive days above the bleaching threshold (D), and 13 year mean *chl-a* concentration from 2002-2015 (E). Maps generated from means calculated from daily satellite measurements taken from Jan 2003-Dec 2012.

**S3 Fig. Linear regression of Physical Parameters vs. NMDS1 and NMDS2 by site type**

Linear regression of average annual max temp (A, F), average annual temp range (B, G), average annual days above the bleaching threshold (C, H), average annual consecutive days above the bleaching threshold (D, I), and *Chl-a* (E, J) vs. NMDS1 and NMDS2 by site type. R^2^ values are included for each regression that yielded a significant slope *(p* <0.05).

**S1 Table. Site locations**

Summary of survey sites, how they were classified, and where they were located (latitude/ longitude).

**S2 Table. *p*-values and R^2^ from Linear Regression of Physical Parameters vs. NMDS1 and NMDS2**

Summary of *p* and R^2^ values for physical parameters vs. NMDS1 and NMDS 2. Significant *p*-values are in bold.

**S3 Table. *p*-values and R^2^ for Linear Regression of Physical Parameters vs. NMDS1 by Site Type**

Summary of *p* and R^2^ values for physical parameters vs. NMDS1 by site type. Significantp- values are in bold.

**S4 Table. Population of major towns in Belize**

Populations of major towns in Belize from 2010-2015. Data source: Statistical Institute of Belize.

## Author Contributions

Planned and designed the field surveying: JHB KDC. Identified sites and designed temperature metric: JHB. Downloaded and manipulated the temperature data FPL. Carried out field work and surveys: JHB JET TAC HEA SWD KDC. Analyzed the data: JHB JET TAC SWD FPL. Wrote/ edited the paper: JHB JET TAC HEA SWD FPL KDC.

## Acknowledgements

We thank D. Hoer, L. Speare, and A. Knowlton for laboratory assistance, P. McDaniel for providing GIS expertise, and C. Berger and S. Hackerott for assistance with coding. We also thank NASA JPL for access to MUR SST data used in this paper, NOAA ERDAAP for access to *chl-a* and temperature data, Belize Fisheries Department for issuing permits that has allowed this research to occur, and Garbutt’s Marine for providing local expert guides and boats for field research. The authors declare that no conflict of interests exists.

## References

1 Hughes TP, Baird AH, Bellwood DR, Card M, Connolly SR, et al. (2003) Climate change, human impacts, and the resilience of coral reefs. Science 301: 929–933.

2 Hoegh-Guldberg O, Mumby PJ, Hooten AJ, Steneck RS, Greenfield P, et al. (2007) Coral Reefs under Rapid Climate Change and Ocean Acidification. Science 318: 1737–1742.

3 Frieler K, Meinshausen M, Golly A, Mengel M, Lebek K, et al. (2013) Limiting global warming to 2 °C is unlikely to save most coral reefs. Nature Climate Change 3: 165–170.

4 Donner SD, Skirving WJ, Little CM, Oppenheimer M, Hoegh-Guldberg O (2005) Global assessment of coral bleaching and required rates of adaptation under climate change. Global Change Biology 11: 2251–2265.

5 Chollett I, Mumby PJ, Muller-Karger FE, Hu CM (2012) Physical environments of the Caribbean Sea. Limnology and Oceanography 57: 1233–1244.

6 Gardner TA, Cote IM, Gill JA, Grant A, Watkinson AR (2003) Long-term region-wide declines in Caribbean corals. Science 301: 958–960.

7 Jokiel PL, Coles SL (1990) Response of Hawaiian and other Indo-Pacific reef corals to elevated temperature. Coral Reefs 8: 155–162.

8 D’Croz L, Mate JL, Oke JE (2001) Responses to elevated sea water temperature and UV radiation in the coral Porites lobata from upwelling and non-upwelling environments on the Pacific coast of Panama. Bulletin of Marine Science 69: 203–214.

9 Aronson R, Precht W, Toscano M, Koltes K (2002) The 1998 bleaching event and its aftermath on a coral reef in Belize. Marine Biology 141: 435–447.

10 Wooldridge S, Done T, Berkelmans R, Jones R, Marshall P (2005) Precursors for resilience in coral communities in a warming climate: a belief network approach. Marine Ecology Progress Series 295: 157–169.

11 Donner SD, Knutson TR, Oppenheimer M (2007) Model-based assessment of the role of human-induced climate change in the 2005 Caribbean coral bleaching event. Proceedings of the National Academy of Sciences of the United States of America 104: 5483–5488.

12 van Hooidonk R, Maynard JA, Liu Y, Lee S-K (2015) Downscaled projections of Caribbean coral bleaching that can inform conservation planning. Global Change Biology: n/a-n/a.

13 Soto I, Muller Karger F, Hallock P, Hu C (2011) Sea surface temperature variability in the Florida Keys and its relationship to coral cover. Journal of Marine Biology 2011.

14 Oliver TA, Palumbi SR (2011) Do fluctuating temperature environments elevate coral thermal tolerance? Coral Reefs 30: 429–440.

15 Barshis DJ, Ladner JT, Oliver TA, Seneca FO, Traylor-Knowles N, et al. (2013) Genomic basis for coral resilience to climate change. Proceedings of the National Academy of Sciences 110: 1387–1392.

16 Pineda J, Starczak V, Tarrant A, Blythe J, Davis K, et al. (2013) Two spatial scales in a bleaching event: Corals from the mildest and the most extreme thermal environments escape mortality. Limnology and Oceanography 58: 1531–1545.

17 Castillo KD, Ries JB, Weiss JM, Lima FP (2012) Decline of forereef corals in response to recent warming linked to history of thermal exposure. Nature Climate Change 2: 756–760.

18 Carilli J, Donner SD, Hartmann AC (2012) Historical temperature variability affects coral response to heat stress. Plos ONE 7: e34418.

19 van Woesik R, Houk P, Isechal AL, Idechong JW, Victor S, et al. (2012) Climate-change refugia in the sheltered bays of Palau: analogs of future reefs. Ecology and Evolution 2: 2474–2484.

20 Fine M, Gildor H, Genin A (2013) A coral reef refuge in the Red Sea. Global Change Biology 19: 3640–3647.

21 Lirman D, Fong P (2007) Is proximity to land-based sources of coral stressors an appropriate measure of risk to coral reefs? An example from the Florida Reef Tract. Marine Pollution Bulletin 54: 779–791.

22 Darling ES, Alvarez-Filip L, Oliver TA, McClanahan TR, Côtô IM (2012) Evaluating life-history strategies of reef corals from species traits. Ecology Letters 15: 1378–1386.

23 Grime JP, Pierce S (2012) The evolutionary strategies that shape ecosystems: John Wiley & Sons.

24 Darling ES, McClanahan TR, Côté IM (2013) Life histories predict coral community disassembly under multiple stressors. Global Change Biology 19: 1930–1940.

25 Alvarez-Filip L, Dulvy NK, Côté IM, Watkinson AR, Gill JA (2011) Coral identity underpins architectural complexity on Caribbean reefs. Ecological Applications 21: 2223–2231.

26 Greenstein B, Curran H, Pandolfi J (1998) Shifting ecological baselines and the demise of Acropora cervicornis in the western North Atlantic and Caribbean Province: a Pleistocene perspective. Coral Reefs 17: 249–261.

27 Buglass S, Donner SD, Alemu JB (2016) A study on the recovery of Tobago’s coral reefs following the 2010 mass bleaching event. Marine Pollution Bulletin 104: 198–206.

28 Alvarez-Filip L, Dulvy NK, Gill JA, Côtô IM, Watkinson AR (2009) Flattening of Caribbean coral reefs: region-wide declines in architectural complexity. Proceedings of the Royal Society of London B: Biological Sciences: rspb20090339.

29 Loya Y, Sakai K, Yamazato K, Nakano Y, Sambali H, et al. (2001) Coral bleaching: the winners and the losers. Ecology Letters 4: 122–131.

30 Van Woesik R, Sakai K, Ganase A, Loya Y (2011) Revisiting the winners and the losers a decade after coral bleaching. Mar Ecol Prog Ser 434: 67–76.

31 Green D, Edmunds P, Carpenter R (2008) Increasing relative abundance of Porites astreoides on Caribbean reefs mediated by an overall decline in coral cover. Marine Ecology Progress Series 359: 1–10.

32 Cramer KL, Jackson JB, Angioletti CV, Leonard-Pingel J, Guilderson TP (2012) Anthropogenic mortality on coral reefs in Caribbean Panama predates coral disease and bleaching. Ecology Letters 15: 561–567.

33 Cramer K. Changes in coral communities and reef environments over the past few centuries in Caribbean Panama; 2010. American Geophysical Union, 2000 Florida Ave., N. W. Washington DC 20009 USA.

34 Cramer KL, Leonard-Pingel JS, Rodríguez F, Jackson JB (2015) Molluscan subfossil assemblages reveal the long-term deterioration of coral reef environments in Caribbean Panama. Marine Pollution Bulletin 96: 176–187.

35 Game ET, McDonald-Madden E, Puotinen ML, Possingham HP (2008) Should we protect the strong or the weak? Risk, resilience, and the selection of marine protected areas. Conservation Biology 22: 1619–1629.

36 Chin TM, Vazquez J, Armstrong E (2013) A multi-scale, high-resolution analysis of global sea surface temperature. Algorithm Theoretical Basis Document, Version 1: 13.

37 Ginsburg R, Lang J (2003) Status of coral reefs in the western Atlantic: Results of initial surveys, Atlantic and Gulf Rapid Reef Assessment(AGRRA) program. Atoll Research Bulletin 496.

38 Simons R (2011) ERDDAP-The Environmental Research Division’s Data Access Program.’. Pacific Grove CA: NOAA/NMFS/SWFSC/ERD.

39 Bell P (1992) Eutrophication and coral reefs—some examples in the Great Barrier Reef lagoon. Water Research 26: 553–568.

40 Bell PRF, Elmetri I, Lapointe BE (2014) Evidence of Large-Scale Chronic Eutrophication in the Great Barrier Reef: Quantification of Chlorophyll a Thresholds for Sustaining Coral Reef Communities. AMBIO 43: 361–376.

41 Polonia ARM, Cleary DFR, de Voogd NJ, Renema W, Hoeksema BW, et al. (2015) Habitat and water quality variables as predictors of community composition in an Indonesian coral reef: a multi-taxon study in the Spermonde Archipelago. Science of The Total Environment 537: 139–151.

42 Team RC (2014) R: A language and environment for statistical computing. R Foundation for Statistical Computing, Vienna, Austria, 2012. ISBN 3-900051-07-0.

43 Gittleman JL, Kot M (1990) Adaptation: statistics and a null model for estimating phylogenetic effects. Systematic Biology 39: 227–241.

44 Dale MR, Fortin M-J (2002) Spatial autocorrelation and statistical tests in ecology. Ecoscience: 162–167.

45 Oksanen J, Blanchet FG, Kindt R, Legendre P, Minchin PR, et al. (2013) Package ‘vegan’. R Packag ver 254: 20–28.

46 Cortés J (1990) The coral reefs of Golfo Dulce, Costa Rica: distribution and community structure: Citeseer.

47 Done TJ (1982) Patterns in the distribution of coral communities across the central Great Barrier Reef. Coral Reefs 1: 95–107.

48 McClanahan T, Maina J (2003) Response of coral assemblages to the interaction between natural temperature variation and rare warm-water events. Ecosystems 6: 551–563.

49 Thompson D, Van Woesik R (2009) Corals escape bleaching in regions that recently and historically experienced frequent thermal stress. Proceedings of the Royal Society of London B: Biological Sciences 276: 2893–2901.

50 McClanahan TR, Ateweberhan M, Omukoto J (2008) Long-term changes in coral colony size distributions on Kenyan reefs under different management regimes and across the 1998 bleaching event Marine Biology 153: 755–768.

51 Grottoli AG, Rodrigues LJ, Palardy JE (2006) Heterotrophic plasticity and resilience in bleached corals. Nature 440: 1186–1189.

52 Anthony K (1999) Coral suspension feeding on fine particulate matter. Journal of Experimental Marine Biology and Ecology 232: 85–106.

53 McClanahan TR, Graham NA, Darling ES (2014) Coral reefs in a crystal ball: predicting the future from the vulnerability of corals and reef fishes to multiple stressors. Current Opinion in Environmental Sustainability 7: 59–64.

54 Aronson RB, Macintyre IG, Wapnick CM, O’Neill MW (2004) Phase shifts, alternative states, and the unprecedented convergence of two reef systems. Ecology 85: 1876–1891.

55 Hughes TP, Tanner JE (2000) Recruitment failure, life histories, and long-term decline of Caribbean corals. Ecology 81: 2250–2263.

56 Wild C, Hoegh-Guldberg O, Naumann MS, Colombo-Pallotta MF, Ateweberhan M, et al. (2011) Climate change impedes scleractinian corals as primary reef ecosystem engineers. Marine and Freshwater Research 62: 205–215.

57 Alvarez-Filip L, Carricart-Ganivet JP, Horta-Puga G, Iglesias-Prieto R (2013) Shifts in coral-assemblage composition do not ensure persistence of reef functionality. Scientific reports 3.

58 Van Woesik R, Tomascik T, Blake S (1999) Coral assemblages and physico-chemical characteristics of the Whitsunday Islands: evidence of recent community changes. Marine and Freshwater Research 50: 427–440.

59 West K, Van Woesik R (2001) Spatial and temporal variance of river discharge on Okinawa (Japan): inferring the temporal impact on adjacent coral reefs. Marine Pollution Bulletin 42: 864–872.

60 Fabricius KE (2005) Effects of terrestrial runoff on the ecology of corals and coral reefs: review and synthesis. Marine Pollution Bulletin 50: 125–146.

61 Marubini F, Atkinson MJ (1999) Effects of lowered pH and elevated nitrate on coral calcification. Marine Ecology Progress Series 188: 117–121.

62 Wooldridge S (2009) A new conceptual model for the enhanced release of mucus in symbiotic reef corals during ‘bleaching’ conditions. Marine Ecology Progress Series 396: 145–152.

63 Szmant AM (2002) Nutrient enrichment on coral reefs: is it a major cause of coral reef decline? Estuaries 25: 743–766.

64 Wooldridge SA (2009) Water quality and coral bleaching thresholds: Formalising the linkage for the inshore reefs of the Great Barrier Reef, Australia. Marine Pollution Bulletin 58: 745–751.

65 Vega Thurber RL, Burkepile DE, Fuchs C, Shantz AA, McMinds R, et al. (2014) Chronic nutrient enrichment increases prevalence and severity of coral disease and bleaching. Global Change Biology 20: 544–554.

66 Zaneveld JR, Burkepile DE, Shantz AA, Pritchard CE, McMinds R, et al. (2016) Overfishing and nutrient pollution interact with temperature to disrupt coral reefs down to microbial scales. Nature communications 7.

67 Paris CB, Cherubin LM (2008) River-reef connectivity in the Meso-American Region. Coral Reefs 27: 773–781.

68 Carilli JE, Prouty NG, Hughen KA, Norris RD (2009) Century-scale records of land-based activities recorded in Mesoamerican coral cores. Marine Pollution Bulletin 58: 1835–1842.

69 Carilli JE, Norris RD, Black BA, Walsh SM, McField M (2009) Local stressors reduce coral resilience to bleaching. Plos ONE 4: e6324.

70 Torres JL, Morelock J (2002) Effect of terrigenous sediment influx on coral cover and linear extension rates of three Caribbean massive coral species. Caribbean Journal of Science 38: 222–229.

71 Hunte W, Wittenberg M (1992) Effects of eutrophication and sedimentation on juvenile corals. Marine Biology 114: 625–631.

72 Carrillo L, Johns EM, Smith RH, Lamkin JT, Largier JL (2015) Pathways and Hydrography in the Mesoamerican Barrier Reef System Part 1: Circulation. Continental Shelf Research 109: 164–176.

73 Sheng J, Tang L (2003) A numerical study of circulation in the western Caribbean Sea. Journal of Physical Oceanography 33: 2049–2069.

74 Tang L, Sheng J, Hatcher BG, Sale PF (2006) Numerical study of circulation, dispersion, and hydrodynamic connectivity of surface waters on the Belize shelf. Journal of Geophysical Research: Oceans 111.

75 Sheng J, Tang L (2004) A two-way nested-grid ocean-circulation model for the Meso-American Barrier Reef System. Ocean Dynamics 54: 232–242.

76 Paris CB, Chérubin LM, Cowen RK (2007) Surfing, spinning, or diving from reef to reef: effects on population connectivity. Marine Ecology Progress Series 347: 285–300.

77 Andrefouet S, Mumby PJ, Mcfield M, Hu C, Muller-Karger RE (2002) Revisiting coral reef connectivity. Coral Reefs 21: 43–48.

78 Prouty N, Hughen K, Carilli J (2008) Geochemical signature of land-based activities in Caribbean coral surface samples. Coral Reefs 27: 727–742.

79 Cooper TF, Uthicke S, Humphrey C, Fabricius KE (2007) Gradients in water column nutrients, sediment parameters, irradiance and coral reef development in the Whitsunday Region, central Great Barrier Reef. Estuarine, Coastal and Shelf Science 74: 458–470.

80 Brown BE (1997) Coral bleaching: causes and consequences. Coral Reefs 16 suppl: s129-s138.

81 Jackson JB, Kirby MX, Berger WH, Bjorndal KA, Botsford LW, et al. (2001) Historical overfishing and the recent collapse of coastal ecosystems. Science 293: 629–637.

82 Burke LM, Maidens J, Spalding M, Kramer P, Green E (2004) Reefs at Risk in the Caribbean: World Resources Institute Washington, DC.

83 Perry C, Larcombe P (2003) Marginal and non-reef-building coral environments. Coral Reefs 22: 427–432.

84 Woesik R, Houk P, Isechal AL, Idechong JW, Victor S, et al. (2012) Climate-change refugia in the sheltered bays of Palau: analogs of future reefs. Ecology and evolution 2: 2474–2484.

85 Graham NAJ, Cinner JE, Norström AV, Nyström M (2014) Coral reefs as novel ecosystems: embracing new futures. Current Opinion in Environmental Sustainability 7: 9–14.

86 Moberg F, Folke C (1999) Ecological goods and services of coral reef ecosystems. Ecological Economics 29: 2151–2233.

